# High-speed atomic force–Raman microscopy

**DOI:** 10.64898/2026.08.06.743386

**Authors:** Keishi Yang, Feng-Yueh Chan, Akihiko Nakamura, Takayuki Uchihashi, Prabhat Verma, Takayuki Umakoshi

## Abstract

A comprehensive understanding of the mechanisms underlying biological systems requires correlative analysis of multiple complementary molecular properties through multidimensional measurements. High-speed atomic force microscopy (HS-AFM) is a powerful tool for elucidating biomolecular structural dynamics at the single-molecule level with high spatiotemporal resolution. However, structural information alone is often insufficient for fully understanding the biological function mechanisms. Here, we report high-speed atomic force–Raman microscopy (HS-AFRM), which enables multimodal measurements combining video-rate structural imaging with chemical-bond analysis. Raman spectroscopy is a powerful, non-invasive technique that probes molecular vibrations to provide chemical information. We achieved several key technical developments that facilitated the seamless integration of HS-AFM and micro-Raman spectroscopy, allowing reliable correlative measurements of structural and chemical information. We validated the versatility of the developed system using representative samples, including two-dimensional materials and a protein. Furthermore, we demonstrate probing of changes in the surrounding environment, which are inaccessible by HS-AFM alone. Multimodal measurements incorporating fluorescence spectroscopy were also demonstrated as an additional practical extension. This multimodal approach substantially enhances the analytical capability of HS-AFM, providing a powerful platform for revealing correlated structural and chemical properties across diverse research fields.

## INTRODUCTION

Multimodal measurement techniques capable of correlatively probing multiple biomolecular properties are essential for a comprehensive understanding of the mechanisms underlying biological function. For elucidating biomolecular structural dynamics, high-speed atomic force microscopy (HS-AFM) is a powerful tool owing to its exceptional spatiotemporal imaging capability under physiological conditions^1^. It provides spatial resolution as high as that of a conventional atomic force microscope (~1 nm), while simultaneously achieving video-rate temporal resolution (>10 frames per second (fps)). This capability enables video-recording of conformational changes in a single protein, contributing to numerous biological discoveries through the direct visualization of dynamic processes such as walking motion of myosin V^2^, rotation of rotor-less F_1_-ATPase^3^, CRISPR-Cas9 DNA editing^4^, and many others^5–14^. More recently, its application has expanded beyond biology to various fields such as materials sciences, polymer chemistry, and organic chemistry, further establishing HS-AFM as a versatile and powerful microscopic tool^15–26^.

In addition, to further enhance its analytical capability, various technical developments have been made in HS-AFM, including stretchable sample substrates and integration with electrochemical measurements^19,25–36^. Among these advances, one of the crucial developments is the stand-alone tip-scan HS-AFM^27,28^. In conventional HS-AFM, the sample stage scans fast in the *x, y*, and *z*-directions. In contrast, in tip-scan HS-AFM, the cantilever performs high-speed scanning on a sample in all three directions. A key advantage of this configuration is that the tip-scan HS-AFM is compatible with an inverted optical microscope as a stand-alone system. Therefore, tip-scan HS-AFM has emerged as a highly promising platform for multimodal measurements, as it can be readily integrated with a variety of optical techniques. Nevertheless, its technical improvements have still been limited to the integration with relatively simple optical or fluorescence microscopy. The combination of advanced optical methods with the high-spatiotemporal-resolution HS-AFM imaging enables the true potential of tip-scan HS-AFM to be fully exploited.

Here, we demonstrate a substantial technical extension of tip-scan HS-AFM by realizing high-speed atomic force–Raman microscopy (HS-AFRM). In this system, micro-Raman spectroscopy is integrated into the tip-scan HS-AFM for correlative analysis of structural and chemical properties. Raman spectroscopy is a widely recognized powerful technique for analyzing molecular vibrations and chemical bonds in a non-invasive and label-free manner by detecting Raman scattered light from samples upon laser irradiation^37^. Moreover, when combined with a tightly focused laser excitation, chemical information can be obtained from the sub-micrometer area. Therefore, micro-Raman spectroscopy has also been a crucial analytical method in the biological fields. By combining this powerful technique with HS-AFM, the analytical capability is significantly enhanced, allowing simultaneous investigation of not only the conformational dynamics of samples but also chemical-bond information. In addition, molecular species that cannot be distinguished solely by morphology can be identified through their Raman signals. Not only samples, but also the surrounding environment such as buffer conditions or the chemical properties of substrates can be characterized, while tracking the structural dynamics of samples. To achieve such comprehensive measurements, we implemented a series of technical developments and constructed an integrated system that enables correlative measurements of micro-Raman spectroscopy and HS-AFM. Using this constructed system, we successfully demonstrated HS-AFRM measurements on several representative samples. HS-AFRM represents a significant advancement, enabling the elucidation of dynamic biological processes with simultaneous structural and chemical insight. Furthermore, the multimodal measurement capability developed here is expected to broaden the applicability of HS-AFM beyond biology into diverse research areas, including materials science, chemical physics, and polymer science.

## RESULTS

### Development of high-speed atomic force–Raman microscopy

Figure 1(a) shows the experimental setup of the constructed HS-AFRM. The lab-built tip-scan HS-AFM was installed on an inverted optical microscope. For excitation of Raman signals, a single-mode laser (wavelength: 532 nm) was used. After passing through several optical components, the laser light was focused on the sample through the oil-immersion objective lens (Olympus, NA 1.45, ×100). The cantilever was also placed at the same area for correlative Raman and HS-AFM measurements. The scattered light from the sample was collected through the same objective, and was detected by a highly sensitive electron-multiplying charge-coupled device (EMCCD) (Teledyne, PROHS-512BX3) after passing through the spectrometer (Teledyne, IsoPlane160). The long-pass edge filter was used to block Rayleigh scattered light. Also, the short-pass filter was inserted to block the HS-AFM laser light used for feedback via an optical lever system. The wavelength of the HS-AFM laser was typically 785 nm, which was changed to 904 nm (Thorlabs, L904P010) to ensure clear separation between HS-AFM laser and Raman signals by the short-pass filter. The acquisition of Raman signals by the EMCCD was triggered by the HS-AFM D/A boards for synchronized measurements, enabling acquisition of a Raman spectrum for every HS-AFM image. In addition, the Raman exposure time can be flexibly extended as needed while maintaining synchronized acquisition with HS-AFM imaging, allowing, for example, one Raman spectrum to be acquired over multiple HS-AFM images.

**Figure 1.**
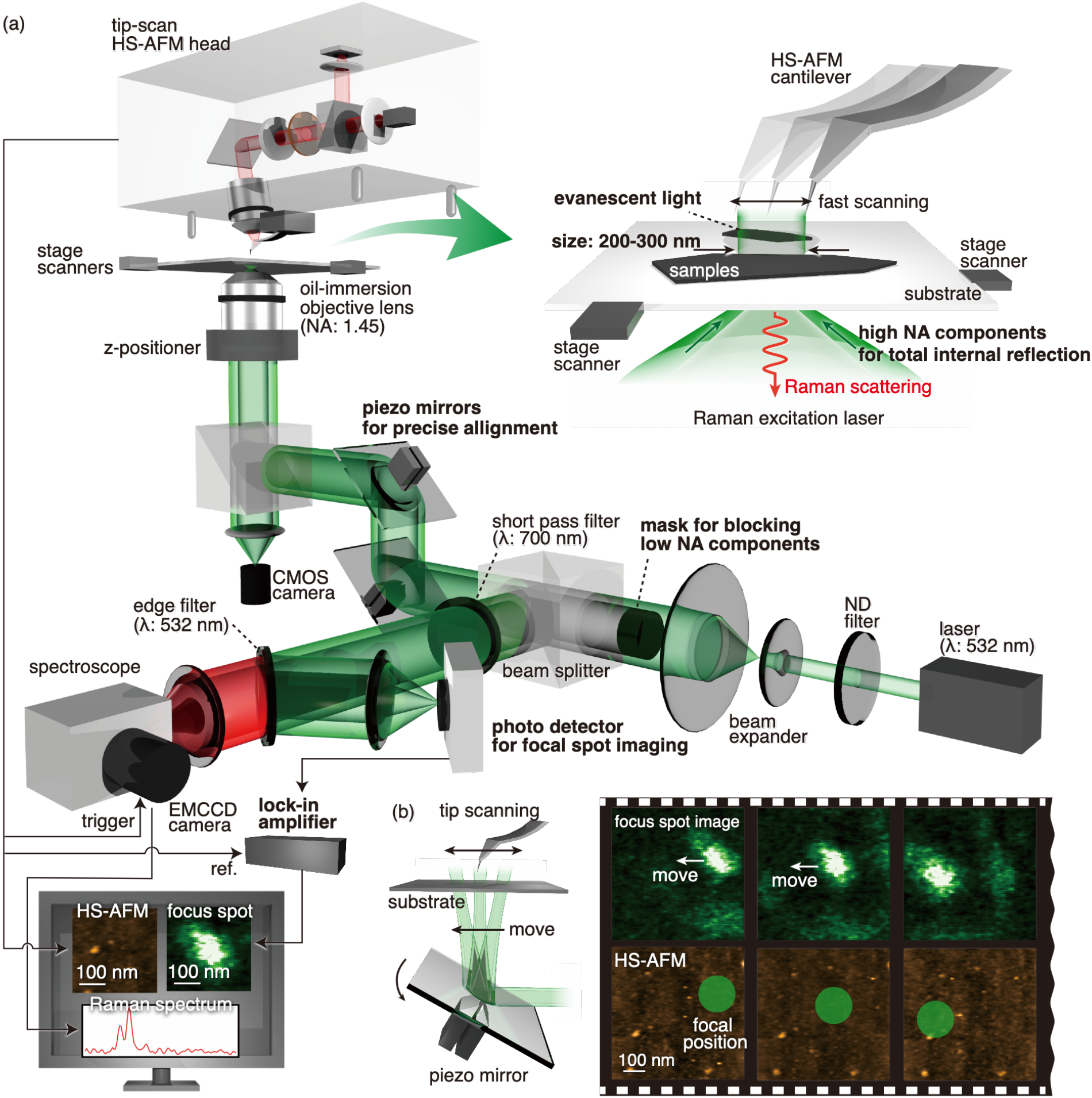
Schematic of the high-speed atomic force–Raman microscope. (a) Overview of the HS-AFRM system. (b) Focal spot imaging and alignment.

Furthermore, to achieve reliable and efficient measurements, we implemented several important technical developments. First, by tightly focusing the excitation light using a high numerical-aperture (NA) oil-immersion objective lens, we reduced the focal spot size to approximately 200–300 nm. Therefore, the focal spot size was almost comparable to the size of the observation area, *i*.*e*., the field of view of HS-AFM, ensuring that Raman signals were collected from almost the same region as that observed by HS-AFM. The positions of the tip and focused laser spot were always aligned, while the observation area was shifted by translating the sample stage using piezo stage scanners. In addition, by introducing a spatial mask to utilize only the high-NA components of the excitation light, samples are excited by an evanescent light generated via total internal reflection illumination, as indicated in the inset of Figure 1(a). Because the evanescent field is localized within approximately 100 nm from the substrate surface, Raman signals are selectively obtained only from the region close to the substrate so that the probing area by Raman measurements is closely correlated to the observation area of HS-AFM. Also, this configuration is essential for largely suppressing Raman signals from samples in the solution as well as unwanted scattered background noise.

A more critical implementation is the method for precisely aligning the tip with the focal laser spot. As the image size of HS-AFM is typically on the order of a few hundred nanometers, the focal spot position needs to be matched to the tip with nanoscale precision. Therefore, we installed custom-built piezo mirrors that enable fine tuning of the focal position to the tip. For positional adjustment, accurate monitoring of the relative positions between the focal spot and the tip is also essential to determine where to finely adjust each position. Simply observing the tip and focal spot using an optical microscope does not provide enough accuracy at the nanoscale. Thus, during HS-AFM imaging, we detected laser light scattered by the tip using a photodetector. When the tip is inside the focal spot, the scattered light intensity increases. By synchronously plotting the scattered intensity together with HS-AFM imaging, an image of the focal spot can be reconstructed, enabling direct visualization of the focal spot position within the HS-AFM field of view. As shown in Figure 1(b) and Supplementary Movie S1, a focal spot image was obtained simultaneously with HS-AFM imaging. In Figure 1(b), a glass coverslip without samples was observed in HS-AFM, and the focal spot was intentionally shifted from right to left by finely tilting the piezo mirrors, confirming precise control of the focal spot within the HS-AFM field of view. In actual HS-AFRM measurements, the focal spot was kept at the center of the HS-AFM imaging area. In addition, because the photodetector detected scattered light reflected at the edge filter, the focal spot observation has no influence on Raman measurements, allowing simultaneous acquisition of all three modalities: focal spot images, HS-AFM images and Raman spectra. In this manner, we established a mechanism that enables real-time visualization of the focal spot position within HS-AFM images with nanoscale precision, while also allowing precise and flexible adjustment via the piezo mirrors during HS-AFRM measurements. In Figure 1(b), to clearly show the focal spot, HS-AFM imaging was performed over a relatively large area (1 × 1 µm^2^). In actual measurements, a smaller imaging area is employed so that the HS-AFM field of view is almost comparable to the focal spot size.

Moreover, the laser light scattered by samples was also detected as background noise in the focal spot image, disturbing the tip–laser alignment. Therefore, the scattered signal was demodulated at the tapping frequency using a lock-in amplifier (Zurich instruments, HF2LI). The cantilever is operated in tapping mode in HS-AFM at a typical oscillation frequency of 300–500 kHz. Scattered light from the tip is modulated at the tapping frequency due to the oscillatory motion, whereas scattered light from the sample is stationary. Thereby, demodulation at the tapping frequency selectively extracts only the tip-scattered signal without any influence from the sample-scattered light. In addition, the use of a lock-in amplifier significantly enhances the detection sensitivity, enabling real-time high-contrast focal spot imaging. Full details of the tip-scan HS-AFM and the integrated HS-AFRM setup are provided in the Supplementary Information Section S1.

### HS-AFM/Raman multimodal measurement

To validate the performance of the developed system, we performed correlative HS-AFM and Raman measurements on two-dimensional (2D) materials, graphene and MoS_2_, as test samples (Figure 2(a)). They are representative 2D materials with excellent optical and electronic properties^38,39^. Both samples were prepared on a glass substrate by mechanical exfoliation using Scotch tapes, where graphene and MoS_2_ were randomly distributed. Two flakes, labeled A and B, were observed in the optical microscope image, as shown in Figure 2(b). In this experiment, the laser focus for Raman measurements was positioned at the center of HS-AFM observation area for simultaneous measurements, as shown in Figure 2(c). While the relative positions of the tip and laser focus were fixed, the sample substrate was translated using the stage piezo scanner to sequentially investigate both flakes. As indicated in Figure 2(b), we started measurements from flake A, and moved to flake B, and returned to flake A. HS-AFM images and Raman spectra were synchronously acquired at rates of 2 fps and 2 spectra per second (sps), respectively. As shown in Figure 2(c) and Supplementary Movie S2, the flat surface morphology of flake A was observed in HS-AFM. Granular particles present on the flake were attributed to residues from the adhesive tapes used for sample preparation. Although we cannot identify the material solely from the HS-AFM image, a characteristic G-band peak at 1580 cm^−1^ was observed in the simultaneously obtained Raman spectrum, which confirmed that flake A was graphene^39^. The sample stage was then moved from flake A toward flake B. At the edge of flake A, the G-band intensity gradually decreased at 11.0 s, and no Raman signal was detected on the bare substrate between flakes A and B, as no sample was present. When the tip reached close to flake B at 55.5 s, the edge of flake B was observed in the HS-AFM image. Concurrently, characteristic Raman peaks of the E^1^_2g_ (386 cm^−1^) and A_1g_ (411 cm^−1^) modes of MoS_2_ were detected, confirming flake B as MoS_2_^38,40,41^. Once the tip moved fully onto flake B at 65.0 s, the Raman signals reached their maximum intensities. The sample stage was subsequently moved back to flake A, where both the graphene morphology and its Raman spectrum were again observed at 115.5 s. These results demonstrate successful correlative measurements combining HS-AFM imaging and micro-Raman spectroscopy, highlighting its capability for simultaneous structural analysis and chemical identification of multiple samples. Furthermore, we confirmed the measurement rates of up to 10 fps for HS-AFM imaging and 10 sps for Raman measurement were possible in our measurement system, as shown in Figure S3 and Supplementary Movie S3.

**Figure 2.**
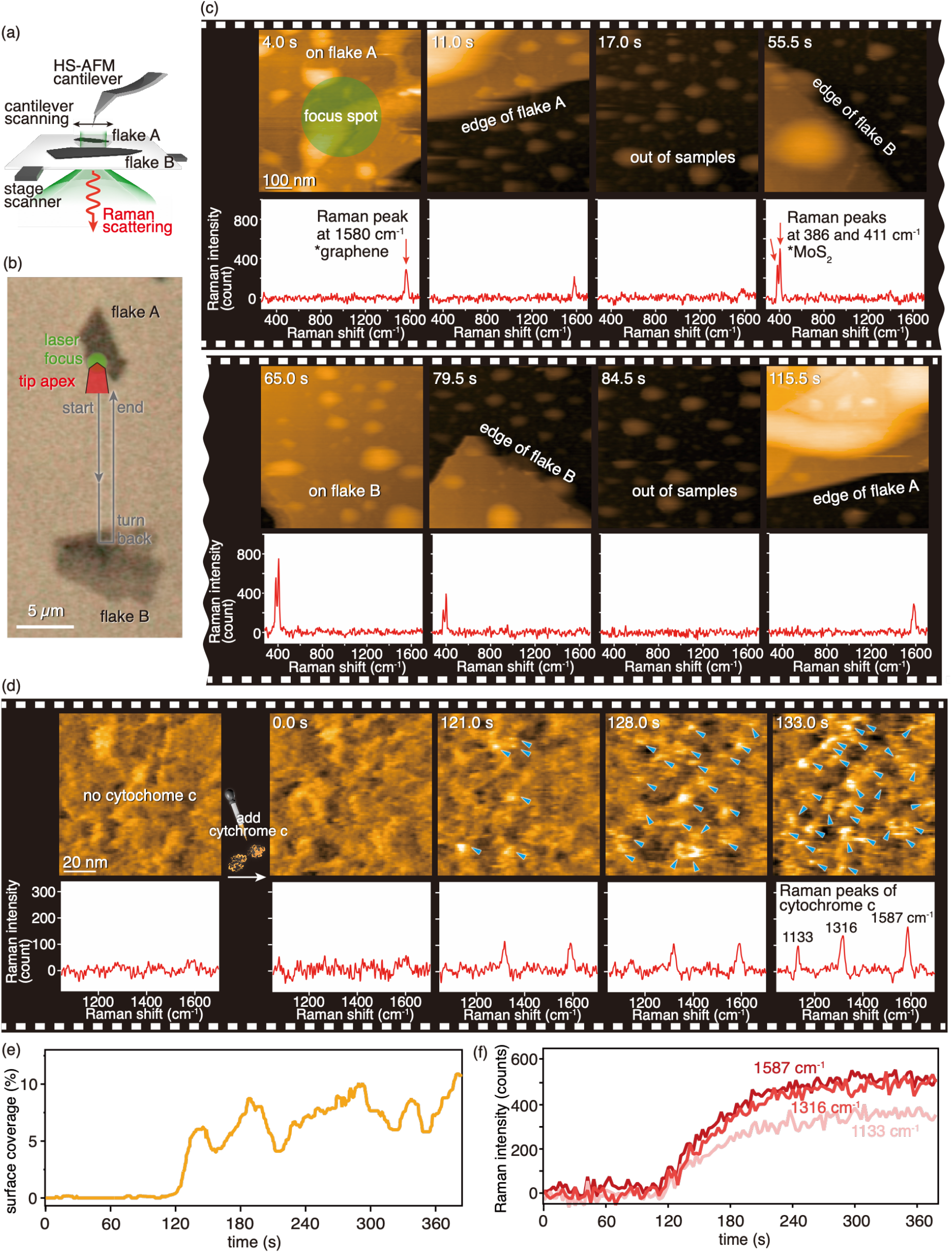
HS-AFRM measurements of 2D materials and biomolecules. (a) Schematic of the HS-AFRM observation of 2D materials. (b) Optical image of flake A and flake B. (c) Snapshots of HS-AFM images and the corresponding Raman spectra acquired simultaneously for the 2D materials. (d) Snapshots of HS-AFM images and the corresponding Raman spectra acquired simultaneously during the cytochrome c adsorption process. (e) Temporal evolution of the apparent surface coverage of cytochrome c. (f) Temporal evolution of representative Raman peak intensities of cytochrome c.

### HS-AFRM measurement of biomolecules

Having validated the correlative accuracy of the developed system using 2D materials, we next applied HS-AFRM for a biological molecule, cytochrome c. Cytochrome c is a well-known heme protein that plays a central role in mitochondrial electron transport^29,42–44^, and has been extensively studied using various spectroscopic techniques, including Raman spectroscopy^42–44^. Here, we observed the adsorption process of cytochrome c on a glass substrate. In the measurement, the substrate surface was functionalized with 3-aminopropyltriethoxysilane (APTES) to render the substrate surface slightly positively charged. As cytochrome c is intrinsically negatively charged, it gently adsorbs onto the APTES-functionalized substrate surface. HS-AFM imaging and Raman measurements were performed at rates of 2 fps and 0.33 sps, respectively, corresponding to the acquisition of one Raman spectrum for every 6 HS-AFM images, thereby allowing longer exposure times in Raman measurements. As shown in Figure 2(d) and Supplementary Movie S4, prior to the introduction of cytochrome c, the substrate surface was observed in sodium phosphate buffer solution, and no Raman peaks were detected. 40 µL of cytochrome c solution was injected into the buffer at 0 s (final concentration: 16 µM). In the beginning, cytochrome c molecules remained dispersed in the solution, so no features were observed in HS-AFM images and Raman spectra. After approximately 100 s, some cytochrome c molecules reached the substrate surface and diffused, as indicated by the blue markers in the HS-AFM images. Simultaneously, characteristic Raman peaks of cytochrome c gradually emerged, including the C-H vibrational mode at 1133 cm^−1^, the A_2g_ mode at 1316 cm^−1^, and the low-spin-state marker peak of the C=C bond at 1587 cm^−1 42–44^. As the adsorption process progressed, both the number of adsorbed molecules in HS-AFM images and the corresponding Raman peak intensities increased steadily. To quantitatively evaluate the adsorption process, the apparent surface coverage of cytochrome c was calculated from the HS-AFM images and compared with the temporal evolution of the Raman peak intensities, as shown in Figures 2(e,f). The apparent surface coverage was calculated by applying a height threshold of 1 nm, where pixels above the threshold were regarded as cytochrome c-covered areas and those below as the substrate. As cytochrome c molecules were first observed in the HS-AFM images, characteristic Raman signals simultaneously emerged in the spectra, confirming a strong correlation between the HS-AFM and Raman measurements. We successfully demonstrated the correlative measurements on biological samples, showing the compatibility of the developed system with biological samples. It should be noted that, as demonstrated in the measurements (0.5 s per frame for HS-AFM imaging and 3 s per spectrum for Raman acquisition), the exposure time for Raman measurements can be flexibly adjusted while maintaining synchronization with HS-AFM imaging. This flexibility enables longer Raman exposure times to compensate for intrinsically weak Raman signals.

### Sensing of surrounding environments enabled by HS-AFRM measurements

We have thus far demonstrated the capability of the developed system for analyzing samples. Moreover, we show that the integration of Raman spectroscopy enables simultaneous probing of changes in the environments surrounding samples. Surrounding environments, such as solution conditions or substrate properties, are essential for biological functions, which however generally cannot be evaluated using HS-AFM. By integrating Raman spectroscopy with HS-AFM, it becomes possible to perform HS-AFM imaging while simultaneously analyzing such environmental properties.

Here, as a simple demonstration, we probed the change in solution conditions between water (H_2_O) and heavy water (D_2_O) in the HS-AFM observation area, as illustrated in the experimental schematic in Figure 3(a). No sample was placed on the glass substrate to simplify the interpretation of the experimental results. While we observed the glass substrate surface in 200 µL of H_2_O using HS-AFM and simultaneously detected Raman signals of H_2_O from the same area, we subsequently injected 50 µL of D_2_O to investigate changes in HS-AFRM measurements during the mixing process. The measurement was performed at the acquisition rates of 2 fps for HS-AFM imaging and 2 sps for Raman measurements, as shown in Figure 3(b) and Supplementary Movie S5. At the beginning of the measurement, HS-AFM images showed the glass substrate with a few minor particles, while a clear Raman peak corresponding to H_2_O (~3400 cm^−1^) was observed in the Raman spectrum. After the injection of D_2_O at ~10.0 s, a new Raman peak corresponding to D_2_O (~2500 cm^−1^) immediately appeared, accompanied by a rapid decrease in the Raman intensity of H_2_O. This indicates that D_2_O quickly reached the HS-AFM observation area. No noticeable change was observed in the HS-AFM images, demonstrating that such information is inaccessible by HS-AFM alone but can be obtained by incorporating Raman measurements. Figure 3(c) shows the temporal evolution of the Raman peak intensities of H_2_O and D_2_O. As the mixing process progressed, the D_2_O peak intensity gradually decreased, while the H_2_O peak intensity recovered. By converting the intensity ratio of D_2_O to H_2_O, calibrated from separately measured data (Figure 3(d)), into the D_2_O concentration, the local concentration of D_2_O around the HS-AFM observation area was estimated as shown in Figure 3(e). Immediately after injection, the local D_2_O concentration rapidly increased to ~60%, followed by a gradual decrease to approximately 30% after ~700 s. This behavior can be explained by the fact that D_2_O initially settled and concentrated at the substrate surface because of its higher density, and subsequently was mixed with H_2_O gradually. The D_2_O concentration did not reach the expected equilibrium value (~20%) within the observation period, indicating that complete mixing had not yet been achieved. This demonstration highlights that this system has an ability to quantify the local solvent changes of the HS-AFM observation area via Raman measurements, which are undetectable by HS-AFM alone.

**Figure 3.**
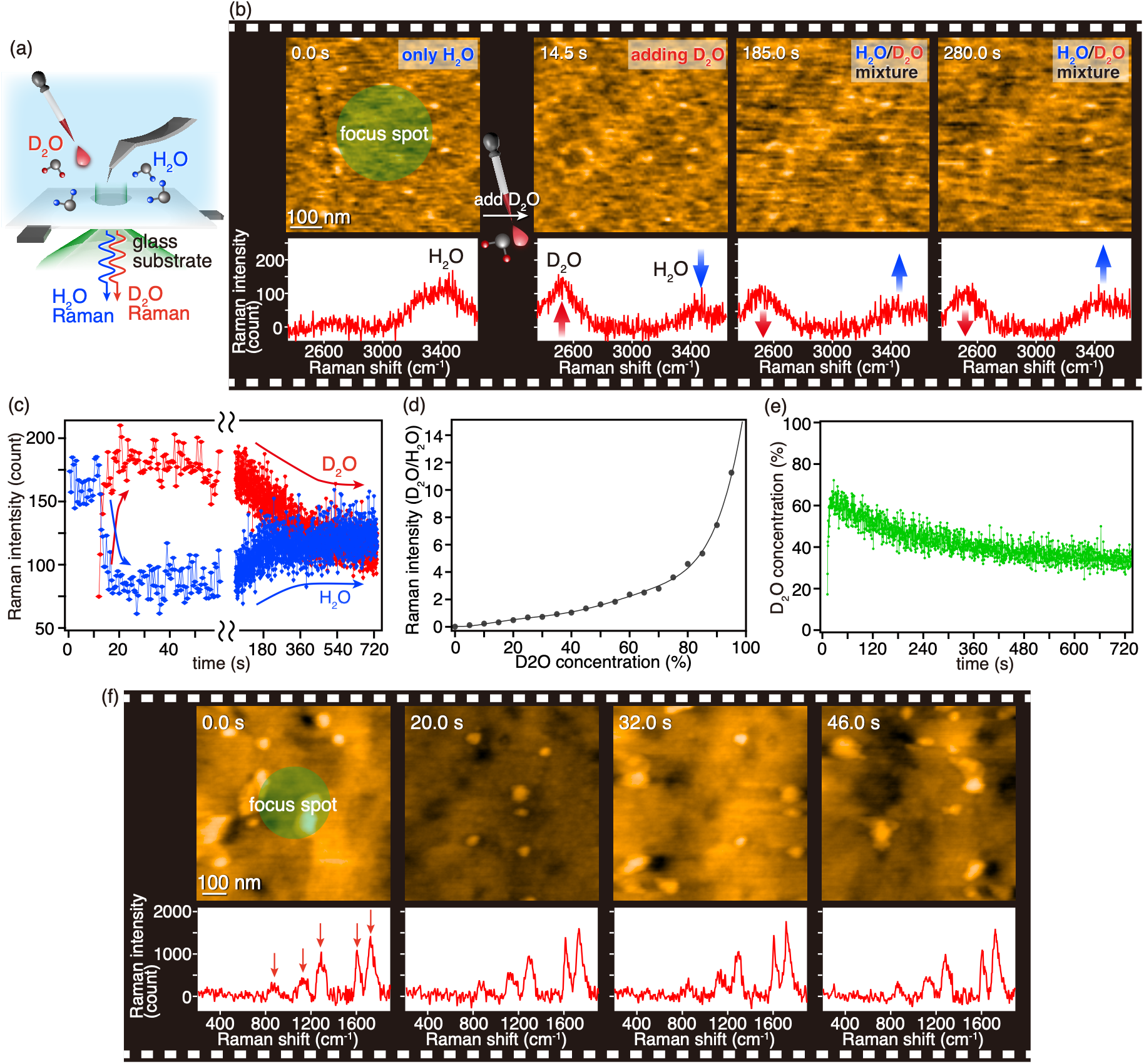
Investigations of surrounding environments enabled by HS-AFRM. (a) Schematic of the HS-AFRM observation of H_2_O/D_2_O mixture. (b) Snapshots of HS-AFM images and the corresponding Raman spectra acquired simultaneously during the mixing process. (c) Temporal evolution of Raman intensities of H_2_O and D_2_O. (d) Relationship between Raman intensity ratio of D_2_O to H_2_O and the concentrations of D_2_O. (e) Temporal evolution of D_2_O concentration, converted from (c,d). (f) Snapshots of HS-AFM images and the corresponding Raman spectra acquired simultaneously for the PET plate.

We further demonstrate that our technique is useful for probing substrate properties. Here, we investigated a polyethylene terephthalate (PET) plate as a representative polymer substrate. We directly placed the PET plate on the sample stage, and performed HS-AFRM measurements. The measurements were conducted at the acquisition rates of 0.5 fps for HS-AFM imaging and 0.5 sps for Raman spectroscopy. As shown in Figure 3(f) and Supplementary Movie S6, HS-AFM images of the PET surface and Raman spectra acquired from the same area were obtained simultaneously. We observed different locations on the PET plate by translating the sample stage, and the corresponding HS-AFM images exhibited distinct surface topographies. We simultaneously detected characteristic Raman peaks of PET, as indicated by the arrows, although no obvious changes were observed under the present measurement conditions. The observed Raman peaks correspond to carbonyl stretching mode (1722 cm^−1^), C=C stretching of benzene ring (1606 cm^−1^), C-C and C-O stretching (1289 cm^−1^), and C-H vibrational mode (863 cm^−1^) of PET^45^. PET is a widely used plastic material in daily life, and there is a strong demand for effective PET degradation methods. Although we focused on measurements of the PET plate alone in this experiment, our HS-AFRM platform has the potential to elucidate PET degradation mechanisms. In particular, simultaneous HS-AFM observation of PET-degrading enzymes, such as PETase, together with Raman spectroscopic analysis of PET properties, including crystallinity, could provide valuable insights into enzymatic degradation processes.

### HS-AFM/fluorescence spectral multimodal measurement

Because our system is capable of detecting Raman signals, it can readily detect fluorescence signals, which are generally much stronger. Therefore, we also demonstrated fluorescence spectral measurements using the same experimental system, as fluorescence spectroscopy is widely employed in biological research for detecting target molecules through fluorescent labelling. Here, we demonstrated correlative HS-AFM and fluorescence spectral measurements using fluorescent polystyrene beads as a test sample. Carboxylate-modified fluorescent beads (diameter: 40 nm) were immobilized on a glass substrate via APTES modification, following a similar procedure used for cytochrome c measurements. The measurements were conducted at acquisition rates of 2 fps for HS-AFM imaging and 2 sps for fluorescence spectroscopy. Figure 4 and Supplementary Movie S7 show HS-AFM images correlated with the simultaneously acquired fluorescence spectra. When a fluorescent bead appeared in the HS-AFM images, corresponding fluorescent signals appeared in the spectra at around 560 nm. These results demonstrate the versatility of the developed system for both Raman and fluorescence spectroscopic measurements. While Raman spectroscopy enables label-free analysis of molecular vibrations, fluorescence spectroscopy provides highly sensitive and specific molecular identification through labelling of target molecules. The ability to directly correlate fluorescence signals with HS-AFM structural dynamics substantially broadens the practical utility of the developed platform, particularly for biological applications requiring identification of specific biomolecules or detection of small molecules unobservable by HS-AFM alone.

**Figure 4.**
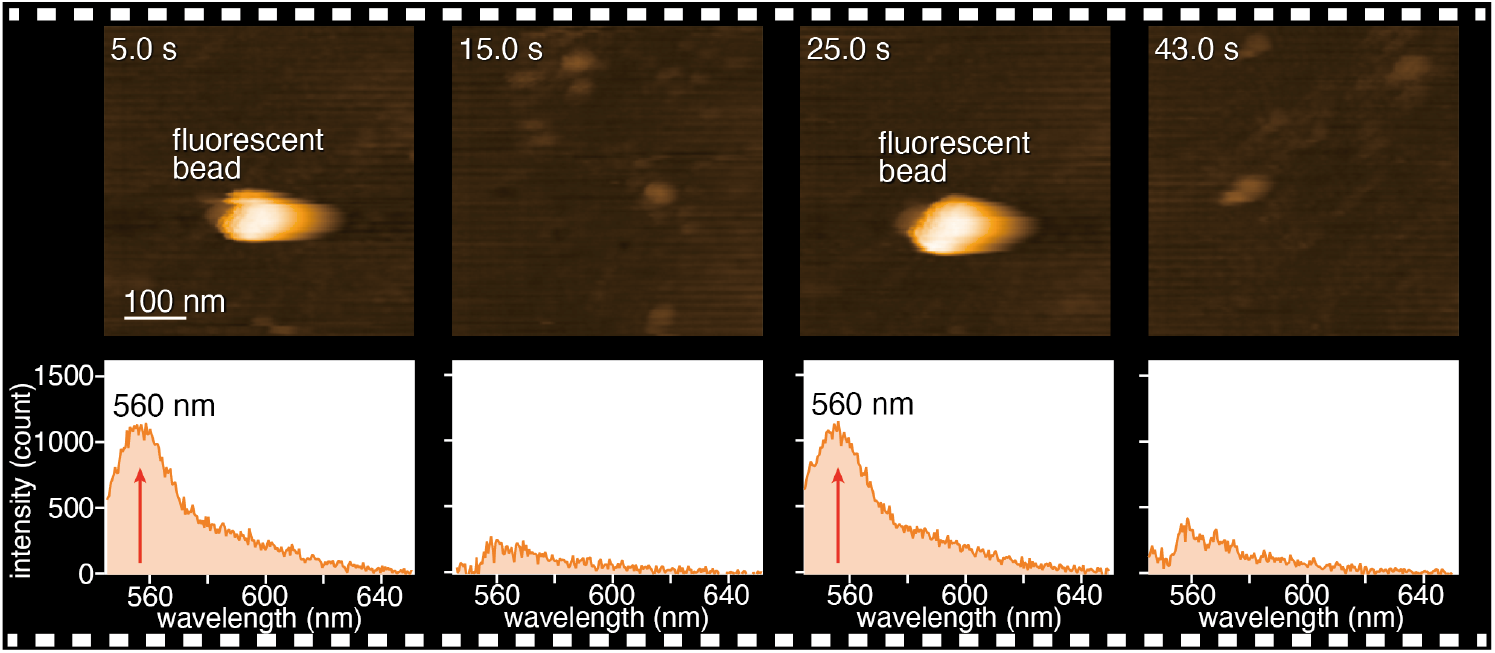
HS-AFM/fluorescence spectral multimodal measurement. Snapshots of HS-AFM images and the corresponding fluorescence spectra acquired simultaneously for fluorescent polystyrene beads.

## CONCLUSION

In this study, we developed a multimodal microscopy platform that integrates real-time nanoscale imaging by HS-AFM with chemical analysis by micro-Raman spectroscopy. We demonstrated its significant capabilities and versatility through a series of representative measurements, ranging from multimodal measurements of 2D materials and biological molecules to analyses of local environmental conditions and fluorescence signals. This substantial enhancement in analytical capability enables correlated observation of structural dynamics together with rich optical information, opening new opportunities for elucidating functional mechanisms of biomolecules and contributing to advances in the biological sciences.

Although Raman spectroscopy usually suffers from limited detection sensitivity, further integration of surface-enhanced Raman scattering is expected to significantly improve sensitivity, potentially reaching the single-molecule level. Furthermore, incorporation of tip-enhanced Raman spectroscopy could enable Raman measurements with nanoscale spatial resolution. Moreover, although the present study focuses primarily on biological applications, the capability of correlatively probing structural dynamics together with chemical information is equally valuable in many other research areas. For example, it will enable correlative analysis of laser-induced structural dynamics and local changes in chemical-bonding states in advanced materials. Our HS-AFRM platform will substantially broaden the applicability of HS-AFM beyond biology into materials science, chemical physics, polymer science, and other disciplines.

## Author contributions

Takayuki Umakoshi conceived and designed this project. K.Y. performed all experiments, analyzed the results, and wrote the manuscript. K.Y., FY.C., Takayuki Uchihashi, and Takayuki Umakoshi constructed the setup for experiments. K.Y. and A.N. prepared the samples. Takayuki Umakoshi supervised this research. All authors contributed to the discussion and finalization of the manuscript.

## Acknowledgement

This work was partially supported by JST FOREST (JPMJFR233Z), JSPS Grant-in-Aid for Scientific Research (KAKENHI) (Grant-in-Aid for Scientific Research (B) JP24K01385, Grant-in-Aid for Transformative Research Areas (A) Publicly Offered Research “Chiral materials science pioneered by the helicity of light” JP25H01624, Grant-in-Aid for Transformative Research Areas (A) Publicly Offered Research “Materials science of meso-hierarchy” JP24H01717, Grant-in-Aid for Challenging Research (Exploratory) JP24K21718), and a research grant from the Takahashi Industrial and Economic Research Foundation. K.Y. acknowledges the JSPS for a Research Fellowship for Young Scientists (JP25KJ1700).

