## Supplementary information for "High-speed atomic force–Raman microscopy"

**Supplementary Movie S1**: HS-AFM and focal spot movies obtained on a glass substrate without samples, which is the source movie used for the snapshots shown in Figure 1(b). HS-AFM images were acquired at 2 fps with a scan size of 1000 × 1000 $nm^2$ (150 × 150 pixels), which was cropped to 600 × 600 $nm^2$ for the movie. The focal spot images were obtained synchronously with the HS-AFM images at the same area.

**Supplementary Movie S2**: HS-AFRM movie of 2D materials, which is the source movie used for the snapshots shown in Figure 2(c). HS-AFM images were acquired at 2 fps with a scan size of 800 × 800 $nm^2$ (200 × 200 pixels), which was cropped to 600 × 600 $nm^2$ for the movie. Raman spectra were simultaneously acquired at 2 sps. The laser power was ~280 μW.

**Supplementary Movie S3**: HS-AFRM movie of 2D materials at a higher acquisition rate. HS-AFM images and Raman spectra were acquired at 10 fps and 10 sps, respectively. The laser power was ~280 μW.

**Supplementary Movie S4**: HS-AFRM movie of the cytochrome c adsorption process, which is the source movie used for the snapshots shown in Figure 2(d). Solution of cytochrome c was added to the observation buffer at 0 s. HS-AFM images were acquired at 2 fps with a scan size of 150 × 150 $nm^2$ (150 × 150 pixels), which was cropped to a 100 × 100 $nm^2$ for the movie. Raman spectra were obtained at 0.33 sps. The laser power was ~550 μW.

**Supplementary Movie S5**: HS-AFRM movie during the mixing process of $H_2O$ and $D_2O$. The glass substrate was observed in $H_2O$, and $D_2O$ was added to $H_2O$ at around 10 s. HS-AFM images were acquired at 2 fps with a scan size of 1 × 1 $\mu m^2$ (150 × 150 pixels), which was cropped to 600 × 600 $nm^2$ for the movie. Raman spectra were simultaneously acquired at 2 sps. The laser power was ~500 μW.

**Supplementary Movie S6**: HS-AFRM movie of PET substrate. HS-AFM images were acquired at 0.5 fps with a scan size of 1 × 1 $\mu m^2$ (200 × 200 pixels), which was cropped to 800 × 800 $nm^2$ in the movie. Raman spectra were simultaneously obtained at 0.5 sps. The laser power was ~140 μW.

**Supplementary Movie S7**: HS-AFM/fluorescence spectral multimodal measurement of fluorescent polystyrene beads. HS-AFM images were acquired at 2 fps with a scan size of 500 × 500 $nm^2$ (150 × 150 pixels). Fluorescence spectra were simultaneously acquired at 2 sps. The laser power was ~140 μW.

### S1. Experimental details of HS-AFRM

The experimental setup was described in Figure 1(a) in the main text. Figure S1 shows a photograph of the constructed HS-AFRM. The Raman optical setup was constructed beneath the home-built tip-scan HS-AFM. A laser (wavelength: 532 nm, Cobolt, 0532-04-01-0100-700) was used for excitation of Raman signals. The laser beam passed through an ND filter and was expanded using a 10× beam expander. A custom-built spatial mask was implemented to achieve total internal reflection illumination, thereby minimizing unwanted scattering noise from the HS-AFM tip as well as from samples above the substrate. The incident laser was focused onto the substrate via an oil-immersion objective lens (NA 1.45: Olympus, UPLXAPO100XO) mounted on a z-piezo positioner (THK PRECISION, PFHW2525R-3000U). The lateral position of laser focal spot was controlled using custom-built piezo-mirrors. The scattered light was filtered using a short-pass filter (Cut-off wavelength: 700 nm, Thorlabs, FESH0700) and a long-pass edge filter (Semrock, LP03-532R-25) before entering the spectrometer (TELEDYNE, IsoPlane160-G1). The Raman signals were detected by a high-sensitivity EMCCD (TELEDYNE, PROHS-512BX3-0). The scattered light from the tip apex was simultaneously collected by a photodetector (Thorlabs, PDA36A2) and demodulated by a lock-in amplifier (Zurich Instruments, HF2LI) at the cantilever resonance frequency (300–500 kHz) to monitor the focal spot position relative to the tip.

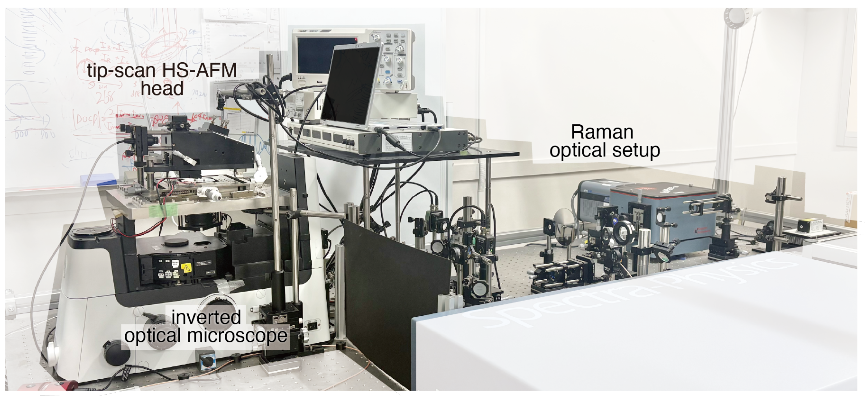


**Figure S1.** Picture of the constructed HS-AFRM.

Figure S2 illustrates a schematic of the optical setup inside the custom-built tip-scan HS-AFM head mounted on an inverted optical microscope (Nikon, ECLIPSE Ti2). A diode laser (wavelength: 904 nm, Thorlabs, L904P010) was employed for optical-lever detection during HS-AFM operation. After passing through a quarter-wave plate, the laser beam was focused onto the cantilever (Olympus, BL-AC10-A2) through an objective lens (NA 0.4: Nikon, CFI160-2) and a glass window. The cantilever, oscillating at its resonance frequency, was raster-scanned over the sample using the tip scanner. To maintain precise laser tracking during HS-AFM operation, a mirror tilter also raster-scans the diode laser synchronously with the scanning motion of the cantilever. This ensures that the laser spot remains focused on the cantilever throughout the scanning. The light reflected from the cantilever was detected by a two-segmented photodetector to generate topographic images, after passing through a polarization beam splitter. A long-pass filter (Thorlabs, FELH0750) was installed to prevent interference from the Raman excitation laser.

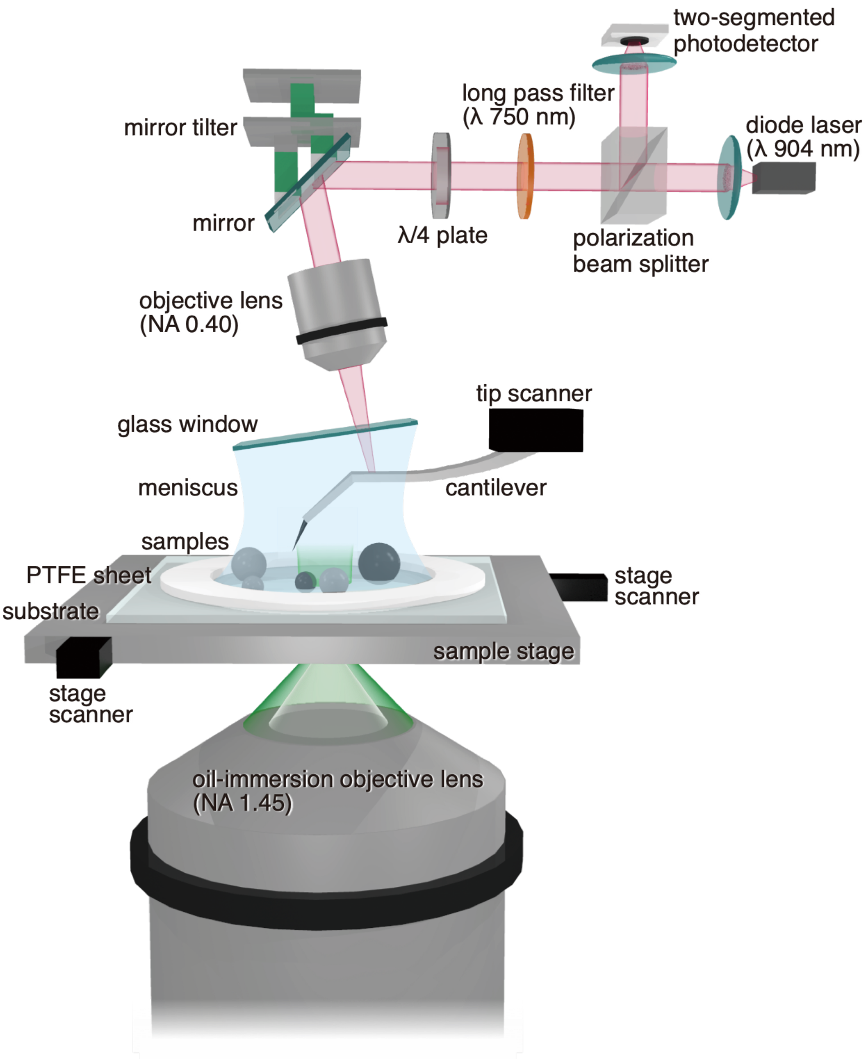


**Figure S2.** Schematic image of the optical setup inside the HS-AFM head.

### S2. Sample preparation

**Substrate for HS-AFRM measurements**

To facilitate simultaneous HS-AFM observation from the top and laser excitation for Raman measurements from the bottom, glass coverslips (22 × 22 $mm^2$, Matsunami, C022221) were used as substrates. The coverslips were cleaned with piranha solution, subsequently rinsed with deionized water, and dried under $N_2$ gas. A donut-shaped PTFE sheet (thickness: 300 μm, inner diameter: ~5 mm, outer diameter: ~20 mm) was glued onto the coverslip as a well to retain solution on the coverslips. Samples were placed on the coverslip inside the center hole of the PTFE sheet. The substrate was mounted on the sample stage and secured with small magnets. ~200 μL of solution was added to form a meniscus between the coverslip and the glass window to ensure stable optical access for the diode laser, as illustrated in Figure S2. Sample solution can be injected from the side of the meniscus during measurements.

**2D materials**

Graphene and $MoS_2$ flakes were prepared via mechanical exfoliation. Initially, the bulk materials were exfoliated using adhesive tape to obtain thin flakes. These flakes were then transferred onto a thermal release tape, which was subsequently pressed against a piranha-cleaned coverslip, before attaching the PTFE sheet. To release the flakes onto the substrate, the assembly was heated to 150℃ using a hotplate (AS ONE, CHPS-170DF). Following the transfer, a donut-shaped PTFE sheet was glued onto the coverslip. Measurements were conducted in deionized water.

**Cytochrome c**

Cytochrome c from equine heart was purchased from Sigma-Aldrich (C2506). Prior to measurement, the surface of the piranha-cleaned substrate was functionalized with 3-aminopropyltriethoxysilane (APTES) to induce a positive surface charge. The substrate was incubated with 5.7 μM APTES for 3 minutes, followed by washing with 1 mL of deionized water. The functionalized coverslip was then mounted on the sample stage and filled with 200 μL of sodium phosphate buffer (10 mM sodium phosphate, pH 7.0, Sigma-Aldrich, 342483). During observation, cytochrome c (100 μM) in buffer solution was gently added to the meniscus to reach a final concentration of 16 μM.

**$H_2O/D_2O$ mixture**

Initial HS-AFM observations were performed on a piranha-cleaned glass substrate in 200 μL of $H_2O$ (deionized water). Subsequently, 50 μL of $D_2O$ (Sigma-Aldrich, 151882) was gently injected into the meniscus to monitor the local solution mixing process.

**PET plate**

Polyethylene terephthalate (PET) film (thickness: 0.25 mm, amorphous, GoodFellow, ES30-FM-000145) was cut into around 23 × 23 $mm^2$ squares and cleaned using a seesaw-type shaker (TAITEC, 0054334-000). For cleaning, the PET film was immersed in a mixture of 80% water and 20% ethanol and shaken for 1 hour at 50 rpm. After cleaning, the film was rinsed with water and dried under $N_2$ gas. To facilitate measurements in solution, a water-repellent coating (Fluoro Surf, FLUORO TECHNOLOGY, FS-1610) was applied to the PET film in a donut-shape. The hydrophobic nature of the Fluoro Surf coating allowed the stable formation of a meniscus between the PET film and the glass window. The PET plate was directly located on the sample stage, and measurements were conducted in deionized water.

**Fluorescent beads**

The piranha-cleaned glass substrate was functionalized by incubating it with 5.7 mM APTES for 3 minutes, followed by washing with 3 mL of water. The solution of carboxylate-modified fluorescent polystyrene beads (diameter: 40 nm, Thermo Fisher Scientific, F8792) was diluted to a concentration of $5\times10^{-5}$% (w/v) in deionized water and incubated on the functionalized substrate for 5 minutes. After incubation, excess beads were removed by washing with 1 mL of deionized water, and the fluorescent beads were measured in deionized water.

### S3. HS-AFRM observation of 2D materials at a high acquisition rate

To evaluate the high-speed measurement performance of the developed system, we performed multimodal measurements at higher acquisition rates of 10 fps for HS-AFM and 10 sps for Raman spectroscopy. As shown in the optical image in Figure S3(a), two flakes, labeled C and D, were selected for high-speed HS-AFRM measurements. As shown in Figure S3(b) and Supplementary Movie S3, despite the shorter exposure time in Raman measurements, the characteristic G-band peak at 1580 $cm^{-1}$ was clearly detected, identifying flake C as graphene. Granular tape residues similar to those observed in the Figure 2(c) in the main text were also observed. Upon moving the stage to flake D, the system immediately captured the characteristic $E^1_{2g}$ (386 $cm^{-1}$) and $A_{1g}$ (411 $cm^{-1}$) Raman modes, confirming flake D as $MoS_2$. Acquisition rates of up to 10 fps and 10 sps were achieved with the constructed system.

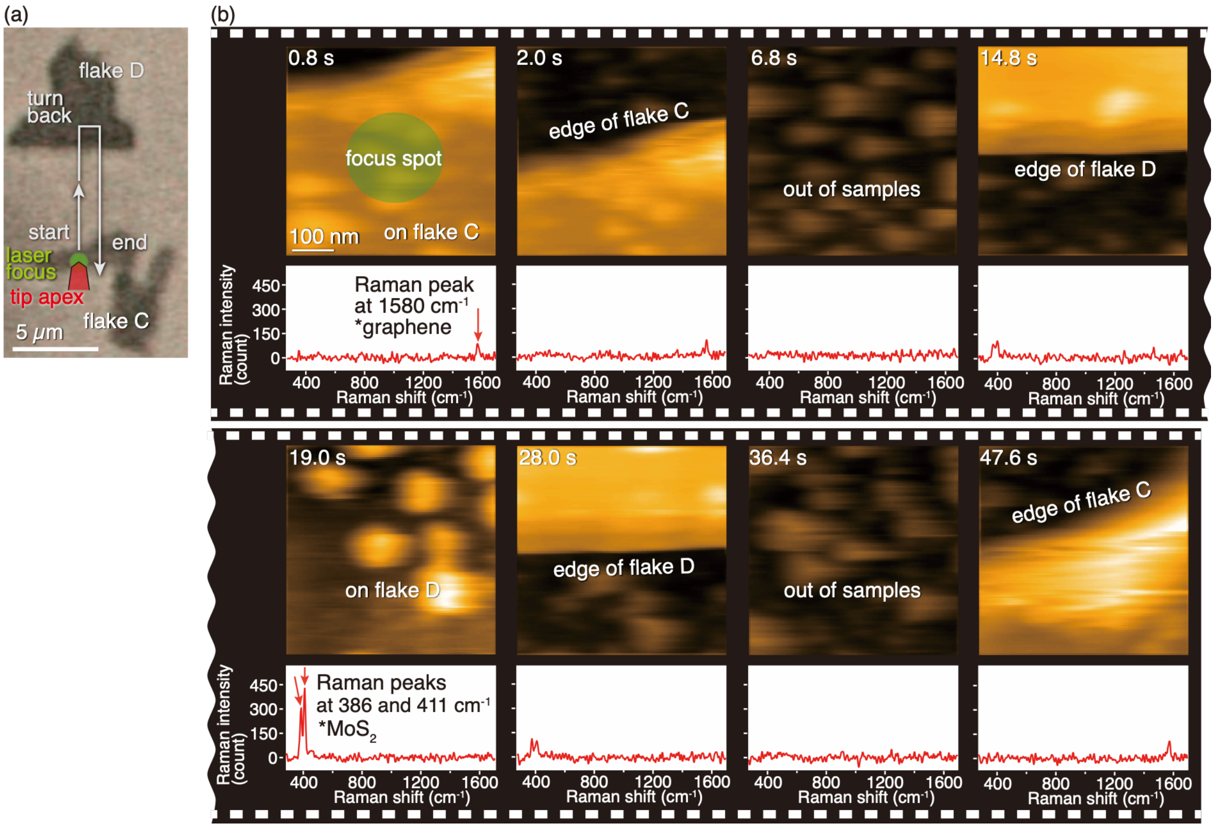


**Figure S3.** (a) Optical image of flake C and flake D. (b) Snapshots of HS-AFRM observation of 2D materials.
